# PSKH1 kinase activity is differentially modulated via allosteric binding of Ca^2+^ sensor proteins

**DOI:** 10.1101/2024.10.20.619163

**Authors:** Christopher R. Horne, Toby A. Dite, Samuel N. Young, Lucy J. Mather, Laura F. Dagley, Jared L. Johnson, Tomer M. Yaron-Barir, Emily M. Huntsman, Leonard A. Daly, Dominic P. Byrne, Antonia L. Cadell, Boaz H. Ng, Jumana Yousef, Dylan H. Multari, Lianju Shen, Luke M. McAloon, Gerard Manning, Mark A. Febbraio, Anthony R. Means, Lewis C. Cantley, Maria C. Tanzer, David R. Croucher, Claire E. Eyers, Patrick A. Eyers, John W. Scott, James M. Murphy

## Abstract

Protein Serine Kinase H1 (PSKH1) was recently identified as a crucial factor in kidney development and is overexpressed in prostate, lung and kidney cancers. However, little is known about PSKH1 regulatory mechanisms, leading to its classification as a “dark” kinase. Here, we used biochemistry and mass spectrometry to define PSKH1’s consensus substrate motif, protein interactors, and how interactors, including Ca^2+^ sensor proteins, promote or suppress activity. Intriguingly, despite the absence of a canonical Calmodulin binding motif, Ca^2+^-Calmodulin activated PSKH1 while, in contrast, the ER-resident Ca^2+^ sensor of the CREC family, Reticulocalbin-3, suppressed PSKH1 catalytic activity. In addition to antagonistic regulation of the PSKH1 kinase domain by Ca^2+^ sensing proteins, we identified UNC119B as a protein interactor that activates PSKH1 via direct engagement of the kinase domain. Our findings identify complementary allosteric mechanisms by which regulatory proteins tune PSKH1’s catalytic activity, and raise the possibility that different Ca^2+^ sensors may act more broadly to tune kinase activities by detecting and decoding extremes of intracellular Ca^2+^ concentrations.

## Introduction

The human kinome was documented more than 20 years ago, yet the attention of the research community has focused on a small subset of kinases that have been disproportionately subjected to detailed study and targeting by small molecule inhibitors. In contrast, approximately one-third of the human kinome has been classified as understudied “dark” kinases^1^, with 22 of these 160 dark kinases residing in the Ca^2+^/Calmodulin-dependent kinase (CAMK) family. Interest in the biological functions of one such CAMK member, Protein Serine Kinase H1 (PSKH1), has recently grown owing to its implication as a crucial factor in kidney development and the association of PSKH1 overexpression with several cancers, including those of the prostate, lung and kidney^2^. Recently, three substitutions in the PSKH1 kinase domain that were identified in patients with kidney ciliopathies were found to suppress PSKH1 catalytic activity^3^, providing a direct link between PSKH1 kinase activity and kidney development.

Among the few studies to date, PSKH1 was reported to reside in the Golgi owing to respective myristoylation and palmitoylation on the conserved residues, Gly2 on Cys3^4^. The precise roles of these post-translational modifications on PSKH1 catalytic activity, however, are not known. Equally, knowledge of PSKH1 interactors and substrates is scant, warranting detailed examination of its interactome and the influence of binders on catalytic regulation. Recent data implicate the Ca^2+^ sensor protein, Calmodulin, as an inhibitor of the activity of another CAMK family member, CHK2^5^, and Calmodulin had been previously reported to inhibit PSKH1 activity^6^. Like PSKH1, CHK2 lacks a conventional Calmodulin binding motif, raising the prospect that PSKH1 may similarly be regulated by Ca^2+^ sensor proteins via an unconventional binding mode. Calmodulin is a ubiquitous intracellular protein in eukaryotic cells, which constitutes up to 0.1% of the total proteome and serves a crucial function as a Ca^2+^ sensor^7^. The cellular influx of Ca^2+^ is a critical second messenger in many facets of biology, from muscle contractility to neuronal communication^7, 8^. Calmodulin binding to four Ca^2+^ ions promotes a conformational change that enables engagement of downstream signaling effectors to regulate their activities through a range of mechanisms, including localization within cells, the occlusion of substrate or partner protein binding, or by modulating a binding partner’s catalytic activity.

Calmodulin is a bilobed, four EF hand fold protein, which typically binds 15-30 amino acid Calmodulin binding motifs in target proteins with up to 20 nM affinity^9, 10^. Our recent data expand the repertoire of Calmodulin binding modes to include regulation of a kinase by direct binding to the kinase domain via a 3D interface on the surface^5^, rather than by binding to a conventional linear sequence motif. Whether Calmodulin regulates PSKH1 activity via a similar mode is yet to be examined. Additionally, PSKH1’s Golgi localization raises the prospect that additional Ca^2+^ sensor proteins in the secretory pathway might contribute to regulation of its catalytic activity^6^. The CREC (Cab45, Reticulocalbin, Erc55, Calumenin) family of Ca^2+^ sensors is known to operate, at least in part, at the Golgi and endoplasmic reticulum^11^, although members are poorly characterized and, unlike Calmodulin, are yet to be implicated in the regulation of kinase activity. Calmodulin is known to bind Ca^2+^ with micromolar affinity to undergo a conformation change that tunes its binding repertoire^10^, and in the presence of a target enzyme, such as MLCK, Calmodulin affinity for Ca^2+^ is vastly elevated to nanomolar range^7, 12^. In contrast, biochemical studies indicate that CREC family sensors undergo an unstructured to structure transition at millimolar Ca^2+^ concentrations^13, 14^, raising the prospect that Calmodulin and CREC proteins may serve complementary roles in Ca^2+^ sensing and kinase regulation in cells. The impact of Calmodulin and CREC proteins on PSKH1 activity therefore is of considerable interest.

Here, we use biochemical and mass spectrometry approaches to define the mechanism of PSKH1 regulation. We identify antagonistic roles for two types of Ca^2+^ sensor, Calmodulin and members of the CREC family, in modulating PSKH1 catalytic activity, via direct binding to the PSKH1 kinase domain. In contrast to the Calmodulin-mediated inhibition of catalytic activity recently observed for CHK2^5^, Calmodulin binding to the PSKH1 catalytic domain promoted enzymatic activity. Another protein identified as proximal to PSKH1 by mass spectrometry, UNC119B, attracted our interest because of its reported contribution to cilia formation^15, 16^, as recently defined for PSKH1^5^. UNC119B is best known as an acyl chain binder^17^, through which activation of Src family kinases and Ras family GTPases has previously been described^18^. Unexpectedly, however, mutation or truncation of the N-terminal myristoylation and palmitoylation sites in PSKH1 did not dampen UNC119B-mediated PSKH1 activation, identifying direct binding to the PSKH1 kinase domain as another mechanism by which UNC119B can promote kinase activity. Our findings illustrate how coordinated interactions with allosteric regulators can complementarily operate to tune kinase catalytic activity. In particular, our data suggest a rheostatic mechanism by which low Ca^2+^ could license Calmodulin to activate kinase activity, while high Ca^2+^ concentrations enable CREC family Ca^2+^ sensors to dampen catalytic activity, to decode extremes in Ca^2+^ flux.

## Results

### PSKH1 substrate consensus motif

As with many understudied kinases, knowledge of PSKH’s physiological substrates and site specificity is scant. To define a consensus substrate motif, we first established methods for robust expression and purification of recombinant PSKH1 from insect cells. Initially, we trialled recombinant wild-type PSKH1 in a radiometric kinase assay with peptide substrates known to be phosphorylated by other CAMK family kinases, and identified the CAMK4 substrate, ADR1, as a target for PSKH1 (**Figure 1A**). We confirmed this activity was attributable to PSKH1 and not a contaminating kinase by comparing the activities of wild-type and D218N kinase-dead mutant PSKH1, with the latter incapable of measurable phosphotransfer (**Figure 1A**).

**Figure 1.**
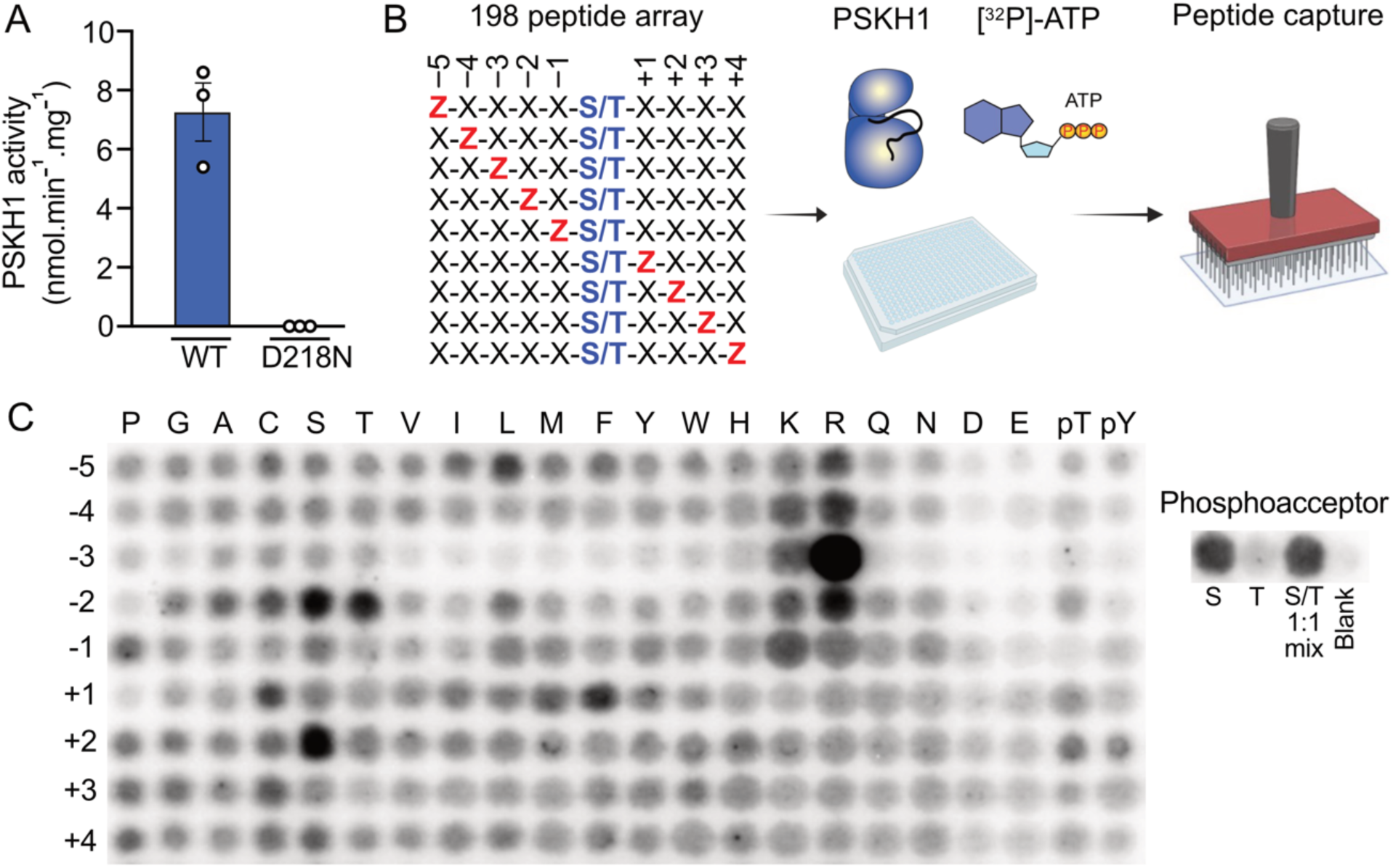
PSKH1 is a serine kinase that favors basic residues at the −3 position. **A** Radiometric assay of recombinant full-length PSKH1 expressed in insect cells. Wild-type, but not D218N kinase-dead, recombinant PSKH1 robustly phosphorylates the ADR1 peptide. **B** Schematic of experimental workflow for positional scanning peptide analysis (PSPA) and representative results; schematic created using BioRender. Z denotes fixed positions containing one of the 20 natural amino acids, or either phosphorylated Thr (pT) or phosphorylated Tyr (pY). X denotes variable positions containing randomized mixtures of all natural amino acids except Ser, Thr and Cys. **C** Radiometric PSKH1 kinase assay performed on peptide array, where darker spots indicate preferred residues. S, G-A-X-X-X-X-X-S-X-X-X-X-A-G-K-K(LC-biotin); T, G-A-X-X-X-X-X-T-X-X-X-X-A-G-K-K(LC-biotin); where X = degenerate mixture of the 16 natural amino acids excluding cysteine, tyrosine, serine, and threonine.

Subsequently, we employed positional scanning peptide array analysis^19, 20, 21, 22^ as an unbiased approach to define the consensus substrate-recognition motif of PSKH1. This method uses an arrayed combinatorial peptide library in which each of nine positions surrounding a central Ser or Thr phosphoacceptor residue are systematically substituted to each of the 20 natural amino acids in addition to phospho-Thr or phospho-Tyr (**Figure 1B**). This approach was recently used to define the substrate preferences of 303 human Ser/Thr kinases^21^, although not all Ser/Thr kinases were able to be profiled in this study, including PSKH1. PSKH1 showed a strong preference for Ser over Thr as the phosphoacceptor residue. This preference is consistent with Leucine occurring as the DFG+1 residue in the PSKH1 activation loop: a hallmark of selectivity towards Serine substrates in kinase families except the CMGC kinases^23^. Like many CAMK group kinases^21^, PSKH1 prefers basic residues N-terminal to the phosphoacceptor residue, with a clear preference for Arg at the −3 position. While Arg is preferred at −3, Lys, but not acetylated or trimethylated Lys, was tolerated (**Figure S1**), emphasizing the required basicity of the −3 residue for PSKH1 recognition. Additionally, while most residues are tolerated at the +1 site, there is an overt preference for Phe, leading to the consensus substrate motif: L/R-X-R-T/R-X-(S)-F-X-X-X where (S) is the phosphoacceptor and X is any amino acid (**Figure 1C**). Serendipitously, the ADR1 peptide sequence, LKKLTRRA(S)FSGG, conforms to the experimentally-derived PSKH1 substrate consensus sequence, which underscores the suitability of ADR1 as a peptide substrate in our radiometric assays. We also noted that the position of the phosphoacceptor could be varied in 7mer peptides, with all but the C-terminal position tolerated, whereas 4mer peptides were poor substrates (**Figure S1**).

### PSKH1 binds secretory network Ca^2+^ sensor proteins and the adaptor, UNC119B

We next sought to identify proteins that bind to, and potentially regulate the activity of, PSKH1 using complementary proteomics approaches (**Figure 2A**). Initially, we C-terminally tagged PSKH1 with the TurboID variant of BirA^24^, to facilitate rapid promiscuous biotinylation of proteins proximal to PSKH1 upon induction of its expression with doxycycline in HEL cells. HEL cells are a human erythroleukemia line that was selected for TurboID experiments because of their endogenous expression of PSKH1 and PSKH2^25^. Relative to uninduced cells, PSKH1 and secretory network proteins, including GOLGA8R, RCN3 (Reticulocalbin-3) and UNC119B, were most enriched following Streptavidin affinity precipitation, tryptic digest and mass spectrometry analysis (**Figure 2B**). We next validated these interactions using the orthogonal proteomics methods, BiCAP (**Figure 2C**) and immunoprecipitation/mass spectrometry (**Figure 2D**) in HEK293T cells. BiCAP involves N-terminal fusion of PSKH1 with a split Venus, which when dimerized can be immunoprecipitated by GFP Trap enrichment (**Figure 2A**). This approach independently validated UNC119B as a PSKH1 interactor (**Figure 2C**). Immunoprecipitation of C-terminally 3C protease cleavable FLAG-tagged PSKH1 and liberation of binders from M2 resin using protease cleavage identified Ca^2+^ sensor proteins of the CREC (Cab45, Reticulocalbin, Erc55, Calumenin) family (**Figure 2D**) to which RCN3 belongs, pointing to a broader role for CREC family proteins in binding to PSKH1. Furthermore, like the pseudokinase paralog, PSKH2^26^, PSKH1 was observed to bind HSP90 and the kinase co-chaperone, Cdc37 (**Figure 2D**), consistent with earlier findings^27^, collectively suggesting that a conserved mechanism may underlie PSKH1 and PSKH2 stabilization.

**Figure 2.**
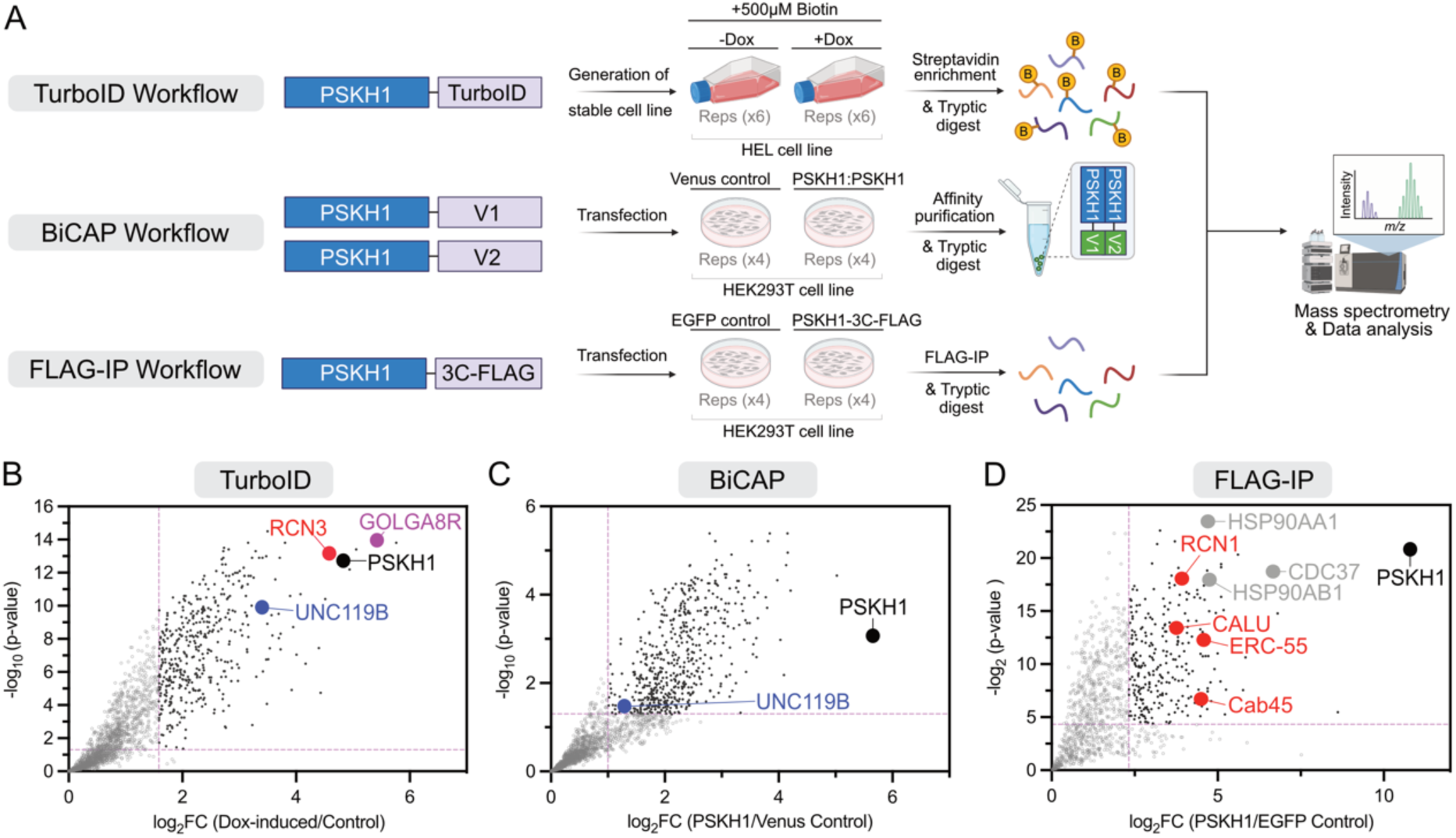
Mapping the PSKH1 interactome using complementary proteomic approaches. **A** Schematic of each experimental workflow; created using BioRender. **B** Volcano plot of TurboID proximity labeling experiment in HEL cells with PSKH1 and proximal proteins of interest highlighted. **C** Volcano plot of BiCAP, validating interaction of dimerized PSKH1 with UNC119B in HEK293T cells. **D** Volcano plot of FLAG-IP, validating interaction of PSKH1 with CREC family members (red dots) and the Cdc37-HSP90 chaperone system (grey). In all panels, PSKH1 is highlighted in black, UNC119B in blue, Calcium sensing interactors of the CREC family in red, secretory pathway interactors in purple and chaperone interactors in grey.

### Calmodulin allosterically activates, and CREC proteins suppress, PSKH1 catalytic output

Recently, we established that Calmodulin directly bound CHK2 – another CAMK family kinase that, like PSKH1, lacks a canonical Calmodulin binding motif – via its kinase domain to suppress CHK2 catalytic activity^5^. While Calmodulin was not observed in our PSKH1 interactome studies, secretory pathway Ca^2+^ sensor proteins were proximal to and bound PSKH1 in HEL and HEK293T cells, respectively, leading us to further investigate the interaction of Ca^2+^ sensor proteins with PSKH1. Firstly, we examined the direct binding of Calmodulin and RCN3, which was observed in our PSKH1 proximitome (**Figure 2B**), to PSKH1 using recombinant proteins in a Far Western assay (**Figure 3A**). These studies verified Calmodulin and RCN3 binding to PSKH1 in a Ca^2+^-dependent manner that was negated by the presence of the Ca^2+^ chelator, EGTA, indicating that a Ca^2+^-mediated conformation change in Calmodulin and a transition to a folded RCN3 structure were crucial to PSKH1 interaction. We next sought to determine the impact of Ca^2+^ sensor protein binding on PSKH1’s catalytic activity in our radiometric assay. Wild-type PSKH1 exhibited robust catalytic activity as measured by ADR1 peptide phosphorylation, and this activity was promoted by Calmodulin ∼10-fold and suppressed by RCN3 ∼2-fold (**Figure 3B**). Calmodulin activation of PSKH1 activity was reduced in the presence of RCN3, indicating that the two Ca^2+^ sensors compete as regulators of PSKH1 kinase activity. We then examined whether two additional CREC family members identified as PSKH1 binders in HEK293T PSKH1 immunoprecipitates, RCN1 and Calumenin, impacted PSKH1 kinase activity. Using our radiometric assay, RCN1 suppressed PSKH1 catalytic activity to a similar extent as RCN3 and also exhibited a reduction in Calmodulin activation when combined, while Calumenin showed no such effect (**Figure 3C**). This suggests that RCN1 and RCN3 bind and inhibit PSKH1 via common regulatory mechanisms.

**Figure 3.**
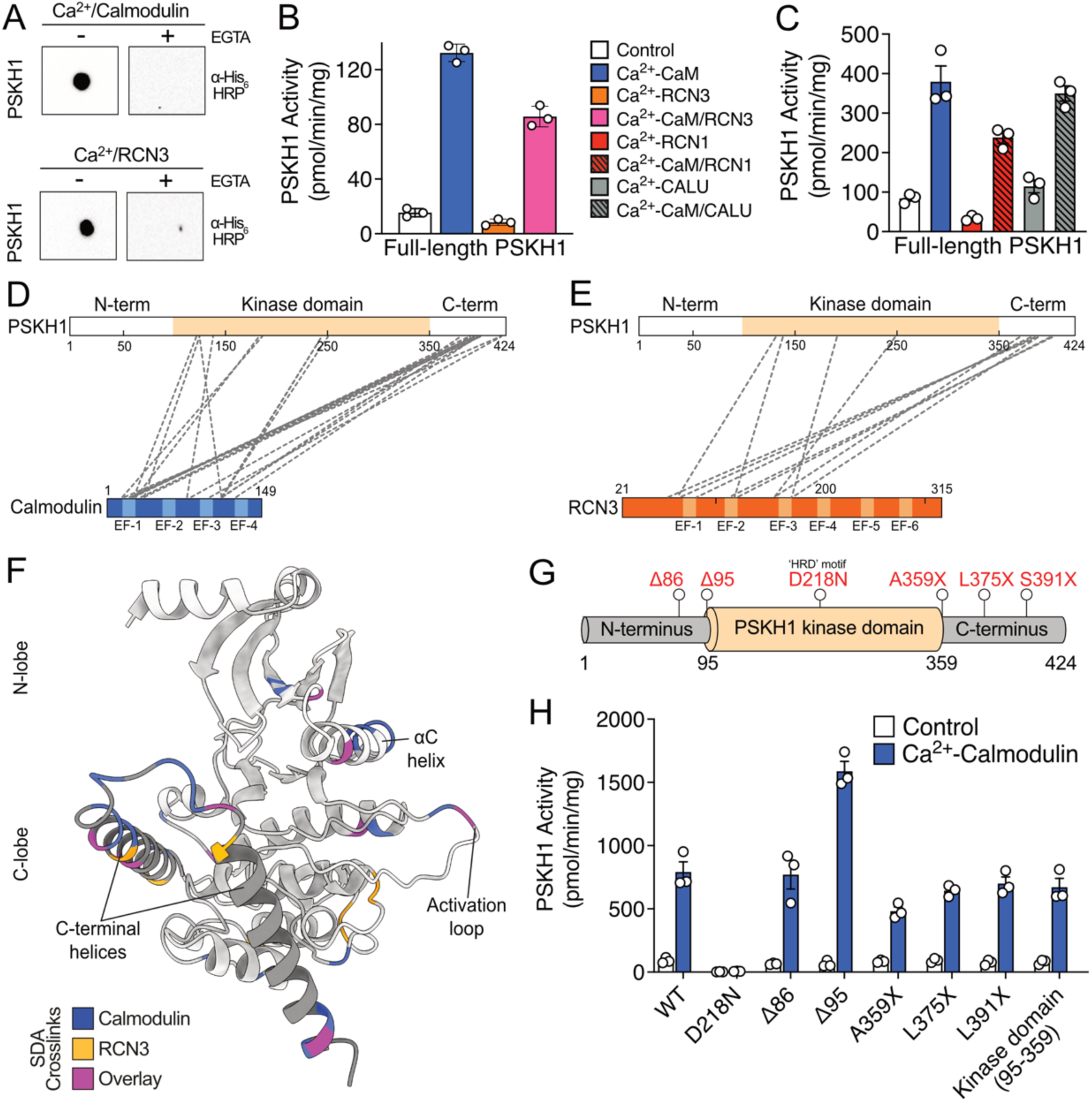
Calcium sensor proteins bind and regulate PSKH1 activity. **A** Far-western blots show that recombinant PSKH1 interacts with His_6_-Calmodulin (CaM) or His_6_-RCN3 in a Ca^2+^-dependent manner (Ca^2+^: 500 μM), as binding is abolished in the presence of the Ca^2+^-chelator, EGTA (10 mM). The interaction was probed using anti-His_6_ HRP antibody. Data are representative of two independent replicates. **B-C** Calmodulin promotes, and Reticulocalbins antagonize, kinase activity in a radiometric assay of HA-tagged PSKH1. **D-E** SDA chemical crosslinking mass spectrometry of PSKH1 with Calmodulin (D) and RCN3 (E). PSKH1 kinase domain is highlighted in beige, Calmodulin in Blue, RCN3 in orange and crosslinks as grey dashes. The position of each EF-hand motif is annotated on Calmodulin and RCN3. **F** AlphaFold model of PSKH1. PSKH1 kinase domain is colored grey and the C-terminal flanking helices in dark grey. **G** Domain architecture of PSKH1, where individual mutations are annotated in red. N-terminal truncations are denoted by the number of residues deleted (Δ) and C-terminal truncations by the position of the introduced stop (X). **H** Radiometric assay of HA-tagged PSKH1 WT and various truncation mutants in the presence and absence of Ca^2+^/Calmodulin.

Because PSKH1 lacks a conventional Calmodulin binding motif typified by a hydrophobic anchor residue and 2-3 basic residues, and CREC sensors are yet to be ascribed roles in kinase regulation, we next sought to define the specific binding interfaces with PSKH1 using crosslinking mass spectrometry and the photoactivatable crosslinker, SDA (**Figure 3D-E**). Calmodulin and RCN3 exhibited similar patterns of interaction, with crosslinks to the central kinase domain and the C-terminal flanking region of PSKH1. Unlike the four EF-hand motif protein, Calmodulin, RCN3 is predicted to harbor 6 EF-hand motifs. Of these, the N-terminal 3 EF-hand motifs interacted with PSKH1, even though studies of other Reticulocalbins suggest the 3 C-terminal EF-hand motifs are capable of adopting globular structures in the presence of Ca^2+^ ^28^. Mapping the crosslinks to an AlphaFold model of PSKH1 revealed a bidentate interaction of Calmodulin and RCN3 with the N- and C-lobes of PSKH1, centered on the regulatory αC helix and activation loop (**Figure 3F**). Further, the model predicts that the C-terminal flanking region of PSKH1 form two α-helices that are positioned adjacent to the activation loop and substrate binding pocket, consistent with the observed crosslinks to this region.

To further investigate whether the regions flanking the core kinase domain contribute to Calmodulin-mediated activation of PSKH1 activity, we prepared a series of truncated PSKH1 proteins and examined their amenability to activation by Calmodulin (**Figure 3G**). As expected, the kinase-dead mutant PSKH1, D218N, did not exhibit activity in the presence or absence of Calmodulin, whilst all truncation mutants – including deletion of both N- and C-terminal regions flanking the core kinase domain – were activated to a similar extent upon addition of Calmodulin, except for an even greater activation of the mutant lacking the N-terminal 95 amino acids (**Figure 3H**). These data demonstrate that Calmodulin binding to the core PSKH1 kinase domain is sufficient for activation, which is supported by the spatial arrangement of the Calmodulin crosslinks to the PSKH1 kinase domain in our AlphaFold model (**Figure 3F**). Activation through direct engagement of the PSKH1 kinase domain is distinct from the conventional mechanism by which Calmodulin activates kinases, which relies on Calmodulin sequestration of a sequence C-terminal to the kinase domain to relieve autoinhibition^5, 29, 30, 31^.

### PSKH1 is allosterically activated by UNC119B

We observed PSKH1 proximity to and interaction with the adaptor protein, UNC119B (**Figure 2B-C**). This interaction drew our attention for several reasons: PSKH1 is myristoylated and palmitoylated at its N-terminus^4^ (**Figure 4A**) and UNC119B is known to function as an adaptor for myristoylated cargo proteins^17, 32, 33, 34^ (**Figure 4B**); UNC119B was previously reported to promote activation of other kinases^18^; like PSKH1^5^, UNC119B has been implicated in ciliary organization^15, 16^; and, lastly, UNC119B interacts with PSKH1’s pseudokinase paralog, PSKH2, which exhibits high amino acid identity to PSKH1 within the kinase domain, despite its catalytic inactivity^26, 35^. We first examined whether UNC119B influenced PSKH1 kinase activity in our radiometric assay (**Figure 4C**) revealing, unexpectedly, that UNC119B promoted PSKH1 catalytic activity. Mutation of the predicted binding site of UNC119B on PSKH1, the myristoylated Gly2, the adjacent palmitoylation site, Cys3, or both together, did not block UNC119B activation of PSKH1 and instead led to elevated basal catalytic activity (**Figure 4C**). These assays revealed that UNC119B binds independently of PSKH1 N-terminal lipidation, suggesting a novel mode of allosteric kinase interaction. We next mapped where UNC119B binds PSKH1 using crosslinking mass spectrometry (**Figure 4D-E**), identifying the interaction mediated principally by the UNC119B C-terminal region with the N-lobe of the kinase domain and flanking region C-terminal to the kinase domain of PSKH1. Truncation analysis supported our crosslinking findings (**Figure 4F**): the region N-terminal to the kinase domain could be deleted without compromising PSKH1 activity, whereas deletion of the flanking region C-terminal to the kinase domain suppressed UNC119B activation of PSKH1. Moreover, deletion of the N- and C-terminal flanks of the kinase domain together did not further compromise UNC119B activation of PSKH1. These data indicate that UNC119B binds and activates the kinase domain of PSKH1, with this interaction augmented by the C-terminal flank of the kinase domain.

**Figure 4.**
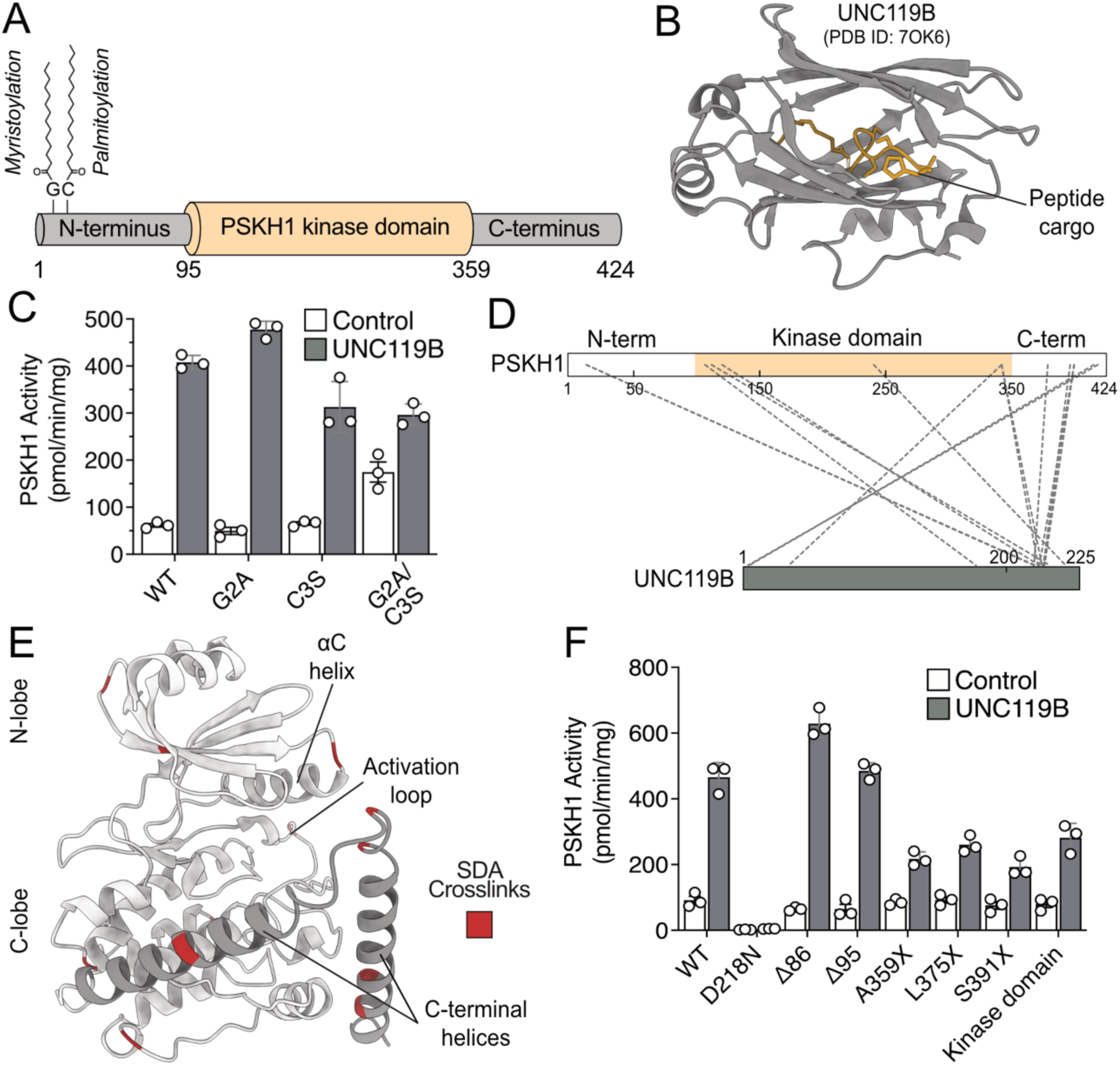
Characterization of the predicted acyl-chain binder, UNC119B. **A** Domain architecture of PSKH1, where the myristoyl and palmitoyl groups at positions 2 and 3 are annotated. **B** Cartoon structure of UNC119B (PDB ID: 7OK6)^34^. The cargo peptide bound in between the β-barrel is colored in gold. **C** Radiometric assay of C-terminal HA-tagged PSKH1 WT, G2A, C3S and a double G2A/C3S mutant. **D** SDA chemical crosslinking mass spectrometry of PSKH1 and UNC119B. PSKH1 kinase domain is highlighted in beige, UNC119B in dark grey and crosslinks as grey dashes. **E** AlphaFold model of PSKH1. SDA crosslinks from (D) are mapped onto the cartoon structure in maroon. The PSKH1 kinase domain is colored grey and the C-terminal flanking helices in dark grey. **F** Radiometric assay of HA-tagged PSKH1 WT and various truncation mutants in the presence and absence of UNC119B. N-terminal truncations are denoted by the number of residues deleted (Δ) and C-terminal truncations by the position of the introduced stop (X), as per Figure 3H.

## Discussion

The biological function of the PSKH1 kinase and the mechanisms underlying its regulation have remained underexplored since initial biochemical studies in the early 2000s^4, 6, 36^. Only recently was the loss of kinase activity causatively linked to kidney ciliopathies in human patients ^3^, supporting an earlier attribution of PSKH1’s involvement in heart cilia organization in mice from a mutagenesis screen for coronary heart disease regulators^37^. A critical challenge to advancing knowledge of PSKH1 biochemistry has been a lack of protocols for producing recombinant protein and the absence of knowledge of substrates or interacting proteins. Here, we overcame these limitations by establishing PSKH1 expression and purification protocols, a robust radiometric assay to measure catalytic activity, defining the PSKH1 interactome using three complementary methods, and detailing PSKH1’s allosteric regulation by Ca^2+^ sensor proteins and the UNC119B adaptor.

An unexpected finding from our study is the prevalence of Ca^2+^ sensor proteins as interactors and regulators of PSKH1 activity. PSKH1 is an ER-resident protein kinase by virtue of N-terminal lipidation, which would localize PSKH1 to secretory organelles proximal to the CREC family of Ca^2+^ sensors – Cab45 (SDF4), Reticulocalbin (RCN)-1 and −3, Erc55 (RCN2), Calumenin. The CREC proteins are predicted to be localized to the ER lumen owing to an N-terminal targeting sequence and a C-terminal HDEL motif that is thought to maintain CREC proteins in the ER^11^. However, the precise subcellular locations of PSKH1 and CREC proteins have not been robustly established experimentally, and detailed exploration awaits the development of suitable, specific reagents. Nonetheless, the observation of CREC proteins in different cell lines in distinct PSKH1 proximitome and interactome experiments indicates their proximity in cells, and the prospect of CREC proteins physiologically modulating PSKH1 activity. To our knowledge, CREC sensors have not previously been ascribed functions in kinase regulation, which makes our finding that each could suppress in vitro PSKH1 kinase activity of great interest. Moreover, RCN3 binding to PSKH1 occurred via an interface akin to that of Calmodulin, despite the latter enhancing activity, whereas RCN3 was inhibitory. PSKH1 regulation through binding of Calmodulin and CREC sensors directly to the kinase domain is reminiscent of our recent finding that Calmodulin can suppress CHK2 kinase activity by binding the kinase domain surface via an interface centred in the activation loop^5^.

Our observations expand the repertoire of kinase regulation modes conferred by Calmodulin and other Ca^2+^ sensor proteins (**Figure 5A**). Perhaps accounting for why there is such diversity in the cellular complement of Ca^2+^ sensing proteins, not only are CREC proteins localized to the secretory machinery like PSKH1, but also regulated distinctly to the archetypal sensor, Calmodulin. Calmodulin undergoes a structural transition on Ca^2+^ binding ^10^ with a K_d_ of 0.5-5 μM^7^. CREC proteins undergo an unfolded-to-folded transition on Ca^2+^ binding^13, 14^, and this only occurs at vastly higher Ca^2+^ concentrations, with K_d_ values for Ca^2+^ reported in the range of 0.1-1 mM^38, 39^. Therefore, while Calmodulin’s transition to a PSKH1-binding competent form requires a modest elevation in Ca^2+^ concentration from the basal intracellular concentration of 0.1 μM, substantively higher and ER-localized flux would be required to license CREC proteins to bind and suppress PSKH1 activity. By extension, we propose a rheostatic model in which the activity of PSKH1 is dictated by proximal Ca^2+^ levels (**Figure 5B**). In this model, Calmodulin activates PSKH1 at low Ca^2+^ and CREC sensors suppress PSKH1 activity at high Ca^2+^ thus allowing dynamic modulation of PSKH1 catalytic activity depending on the Ca^2+^ flux into cells or release from intracellular stores.

**Figure 5.**
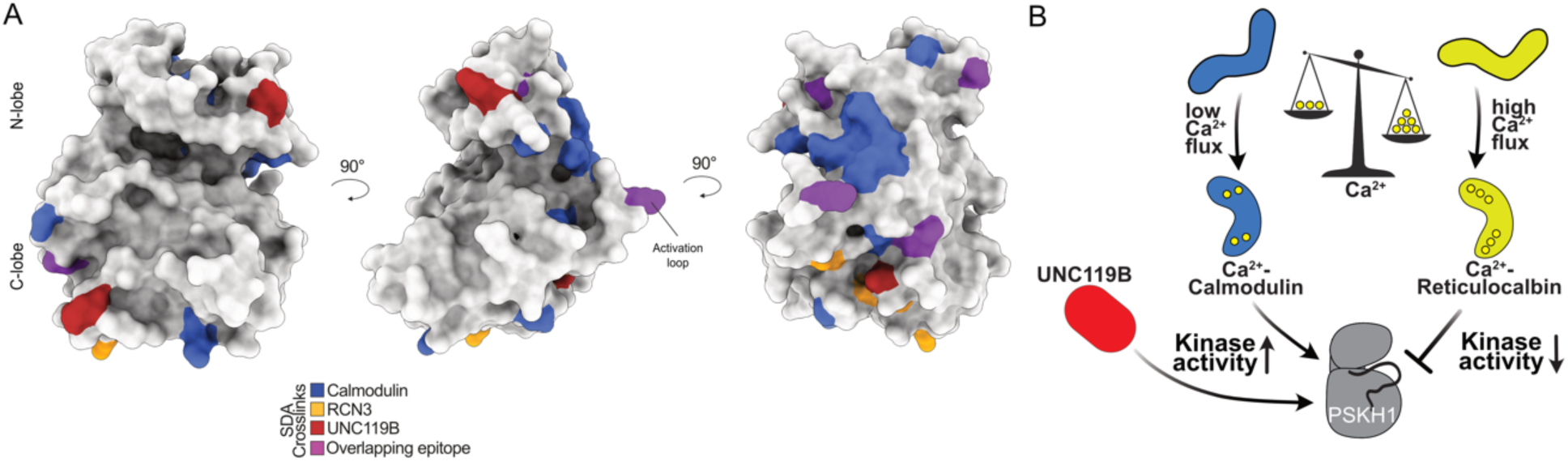
Novel allosteric binders converge on common sites on PSKH1. **A** Orthogonal views of sites bound by Calmodulin (blue), RCN3 (yellow), UNC119B (red) and all interactors (purple). **B** Schematic overview of positive and negative allosteric binding interactions identified in this work.

In addition to Ca^2+^ sensing proteins, our interactomics identified the chaperone protein, UNC119B, which is best described in facilitating protein trafficking between intracellular membranes via its binding to N-myristoylated peptides in its cargoes^32, 34, 40, 41^. An analogous mode of binding to the myristoylated N-terminus of Src family kinases was previously reported to promote kinase activity by relieving lipid occlusion of the kinase active site. Here, we have identified a distinct mode of UNC119B activation of a kinase – where direct binding to the kinase domain, independent of lipidation, promoted kinase activity. Interestingly, UNC119B was identified in our BiCAP mass spectrometry experiments, as an interactor of dimerized PSKH1, raising the possibility that UNC119B binds higher order assemblies of PSKH1 to promote their kinase activity. Interaction of UNC119 with PSKH1, as well as the pseudokinase paralog, PSKH2, has been previously observed in interactome screens^42, 43, 44^, although whether these interactions could serve regulatory functions had not previously been explored. Understanding the precise mode of interaction of UNC119B with PSKH1, and whether this differs from the activating interaction with Calmodulin, awaits high resolution structure determination. However, it is worth noting that crosslinks for each of Calmodulin and UNC119B in our structural mass spectrometry studies map to proximal sites in an AlphaFold model of PSKH1 (**Figure 5**). Such an overlap raises the possibility that a regulatory hotspot may exist on the PSKH1 surface that can be allosterically activated via a shared mechanism, despite the vastly different folds of each activator protein.

Collectively, our study has advanced knowledge of the understudied protein kinase, PSKH1, and identified new mechanisms by which kinases can be allosterically regulated by protein interactors. This study highlights that kinase domains, and not just their flanking regulatory sites, can be targeted at a common site by regulatory proteins of distinct fold to promote catalytic activity. Our findings implicate the understudied CREC Ca^2+^ sensor family as underappreciated protein kinase regulators, and highlight their potential to complement Calmodulin in regulating kinase activity depending on extreme ranges of Ca^2+^ flux. Furthermore, our findings highlight the complexity of mechanisms that operate synergistically to regulate kinase outputs. The applicability of the newly identified modes of Ca^2+^ sensor and UNC119B regulation of PSKH1 to other kinases, including those that remain understudied, awaits careful biochemical examination.

## Materials and Methods

### Expression constructs

The gene coding for full-length human PSKH1 (Uniprot P11801) was synthesised by Gene Universal (Delaware, USA) and subcloned into a mammalian expression vector, pcDNA3.1(-), using the restriction sites XhoI and HindIII with a C-terminal HA tag. PSKH1 variants were introduced into the wild-type template using NEBuilder HiFi DNA Assembly (New England Biolabs). PSKH1 was also subcloned into the insect expression vector, pFastBac GST-GFP, as an in-frame fusion with a TEV protease-cleavable N-terminal GST tag and 3C-protease-cleavable N-terminal GFP tag using the restriction sites BamHI and EcoRI. For TurboID experiments, PSKH1 was subcloned into the mammalian expression vector, pFTRE3G PGK puro, as an in-frame fusion with a C-terminal FLAG tag and TurboID fusion using the restriction sites BamHI and NheI. For BiCAP experiments, PSKH1 was first subcloned into the Gateway Cloning donor vector, pDONR221. The two BiCAP expression vectors were subsequently generated by recombination of the PSKH1 sequence into the pDEST-V1-ORF and pDEST-V2-ORF destination vectors using Gateway Cloning. For FLAG-IP experiments, PSKH1 was subcloned into a pcDNA3 backbone vector as an in-frame fusion with a C-terminal, 3C protease-cleavable FLAG epitope tag using restriction cloning. The gene coding for full-length Calmodulin (CaM; Uniprot P0DP23) was synthesised by IDT (Iowa, USA) as a gBlock and subcloned into the bacterial expression vector, pPROEX Htb (Life Technologies), as an in-frame fusion with a TEV protease-cleavable N-terminal hexahistidine tag using the restriction sites BamHI and NotI. The gene coding for full-length Reticulocalbin-3 (RCN3; Uniprot Q96D15) was synthesised by IDT (Iowa, USA) as a gBlock and subcloned into the bacterial expression vector, pPROEX Htb (Life Technologies), as an in-frame fusion with a TEV protease-cleavable N-terminal hexahistidine tag using the restriction sites BamHI and EcoRI. The gene coding for full-length UNC119B (Uniprot A6NIH7) was synthesised by GenScript (New Jersey, USA) and subcloned into the insect expression vector, pFastBac GST, as an in-frame fusion with a TEV protease-cleavable N-terminal GST tag using the restriction sites BamHI and EcoRI. The gene coding for full-length RCN1 was a kind gift from Naoto Yonezawa (Chiba University)^13^. The gene encoding full-length Calumenin (CALU; Uniprot O43852) was synthesised by GenScript (New Jersey, USA) and subcloned into the bacterial expression vector, pGEX-2T-TEV, as an in-frame fusion with a TEV protease-cleavable N-terminal GST tag using the restriction sites BamHI and EcoRI. Insert sequences were verified by Sanger sequencing (AGRF, VIC, Australia).

### Transient Expression and Immunoprecipitation

Full-length PSKH1 with a C-terminal HA-tag was expressed in HEK293T cells grown in DMEM (Thermo Fisher) media, supplemented with 8% (v/v) Fetal Calf Serum (FCS; Sigma) at 37 °C with 5% CO2. The cells were transfected at 60% confluency using FuGene HD (Roche Applied Science) with 2 µg of pcDNA3.1(-) vector DNA. After 48 h, transfected cells were harvested by rinsing with ice-cold PBS, followed by rapid lysis in situ using lysis buffer (50 mM Tris-HCl pH 7.4, 1% (v/v) Triton X-100, 1 mM PMSF, 1 mM EDTA, 150 mM NaCl, 2 mM Sodium Vanadate, 10 mM NaF) containing a Complete protease inhibitor tablet (Roche). Insoluble debris were removed by centrifugation and supernatants were mixed with 100 µL of anti-HA agarose (50% v/v; Sigma) pre-equilibrated in lysis buffer, followed by successive washes in lysis buffer containing 1 mM NaCl, and finally resuspended in 50 mM HEPES, pH 7.4. Total protein content was quantified using the BCA Protein Assay (Thermo Fisher Scientific), according to manufacturer’s instructions.

### Recombinant Expression and Purification

PSKH1 and UNC119B, both harboring a N-terminal GST tag, were expressed and purified from Expi*Sf*9 insect cells. Briefly, the bacmid was prepared in DH10MultiBac *Escherichia coli* (ATG Biosynthetics) from a pFastBac GST-GFP vector containing PSKH1 as a BamHI-EcoRI insert. *Sf*21 insect cells (Merck) were cultured in Insect-XPRESS (Lonza) media. Each bacmid (1μg) was introduced into 0.9 × 10^6^ *Sf*21 cells by Cellfectin II (Thermo Fisher Scientific) mediated transfection in six-well plates using the Bac-to-Bac protocol (Thermo Fisher Scientific), as detailed elsewhere^45^. After 4 days of static incubation at 27 °C in a humidified incubator, the resulting P1 baculovirus was harvested and added to 50 mL *Sf*21 cells at 0.5 × 10^6^ cells/mL density at 4% v/v, which were shaken at 27 °C, 130 rpm. The cell density was monitored daily using a haemocytometer slide and maintained at 0.5–3.0 × 10^6^ cells/mL by diluting with fresh Insect-XPRESS medium when necessary, until growth arrest (defined as a cell density less than the twice the cell count 1 day prior). Approximately 24 h after growth arrest was recorded, the P2 baculovirus was harvested by collecting the supernatant after pelleting the cells at 500 × *g* for 5 min. P2 baculovirus (3 mL) was added to 0.5 L Expi*Sf*9 cells, seeded at 6 × 10^6^ cells/mL into 2.8 L Fernbach flasks using Expi*Sf* CD Medium (Thermo Fisher Scientific). Expi*Sf*9 cells were harvested 72 h post transduction at 500 ×*g* and pellets snap frozen in liquid N_2_ and stored at −80 °C.

CaM, RCN3, RCN1 and CALU were expressed in *E. coli* BL21-CodonPlus-RIL (Agilent) cells cultured in Super Broth supplemented with ampicillin (100 µg/mL) at 37 °C with shaking at 220 rpm to an OD_600_ of ∼0.6–0.8. Protein expression was induced by the addition of isopropyl β-D-1-thiogalactopyranoside (IPTG; 250 µM) and the temperature was lowered to 18 °C for incubation overnight. Bacterial cells were harvested by centrifugation at 2000 ×*g* and pellets snap frozen in liquid N_2_ and stored at −80 °C.

PSKH1, UNC119B and CALU cell pellets were resuspended in GST buffer (20 mM HEPES pH 7.5, 200 mM NaCl, 10% v/v glycerol), supplemented with cOmplete Protease Inhibitor (Roche) and 0.5 mM Bond-Breaker TCEP [tris-(2-carboxyethyl)phosphine] (ThermoFisher Scientific) and were lysed by sonication, before the lysate was clarified by centrifugation (40,000 ×*g*, 45 min, 4 °C). The proteins were maintained at 4 °C throughout purification. The clarified lysate was incubated with Glutathione Xpure Agarose resin (UBP-Bio) pre-equilibrated in GST buffer for >1 h on rollers at 4 °C, before the beads were pelleted at 500 ×*g* and washed extensively with GST buffer. PSKH1 was further purified by cleaving the GST tag (on resin) with recombinant His_6_-TEV protease overnight at 4 °C. The supernatant was then syringe filtered (0.2 μm), spin concentrated (30 kDa MWCO; Millipore), and then loaded onto a HiLoad 16/160 Superdex 200 pg column (Cytiva) pre-equilibrated with SEC buffer (20 mM HEPES pH 7.5, 200 mM NaCl, 5% v/v glycerol). Purified fractions, as assessed following resolution by reducing StainFree SDS-PAGE gel electrophoresis (Bio-Rad), were pooled, spin concentrated, aliquoted, and snap frozen in liquid N_2_ for storage at −80 °C.

CaM, RCN3 and RCN1 cell pellets were resuspended in Ni-NTA buffer (20 mM HEPES pH 7.5, 200 mM NaCl, 10% v/v glycerol), supplemented with 10 mM imidazole (pH 8.0), EDTA-free cOmplete Protease Inhibitor (Roche) and 0.5 mM Bond-Breaker TCEP and were lysed by sonication, before the lysate was clarified by centrifugation (40,000 × *g*, 45 min, 4 °C). The clarified lysate was incubated with Ni-NTA resin (Roche) pre-equilibrated in Ni-NTA buffer with 5 mM imidazole for >1 h on rollers at 4 °C, before the beads were pelleted at 500 × *g* and washed extensively with Ni-NTA buffer containing 35 mM imidazole. The proteins were eluted in Ni-NTA buffer containing 250 mM imidazole. The eluate was further purified by cleaving the His_6_ tag using recombinant His_6_-TEV protease (for applications where the absence of His_6_ tag was desirable), dialysis, Ni-NTA resin addition to eliminate uncleaved material and the TEV protease. Both cleaved samples were spin concentrated (10 kDa MWCO; Millipore), before being loaded onto a HiLoad 16/160 Superdex 200 pg column (Cytiva) pre-equilibrated with SEC buffer (20 mM HEPES pH 7.5, 200 mM NaCl, 5% v/v glycerol). Purified fractions, as assessed following resolution by reducing StainFree SDS-PAGE gel electrophoresis (Bio-Rad), were pooled and spin concentrated, aliquoted, and snap frozen in liquid N_2_ for storage at −80 °C.

### Generation of TurboID cell lines and validation

Midiprep DNA was co-transfected into HEK293T cells (originally sourced from ATCC) with helper plasmids pVSVg and pCMV δR8.2 to generate lentiviral particles using Effectene (Qiagen). HEL cells (originally sourced from ATCC) were then stably transduced with the resulting lentivirus and successful transductants selected using puromycin (1.25 µg/mL; StemCell Technologies) as before^46, 47^. Transduced HEL cells were cultured in RPMI + 8% FCS, at 37 °C and 10% (v/v) CO_2_ and were routinely PCR monitored for mycoplasma contamination. Expression was validated by Western Blot. Briefly, HEL cells were seeded into 24-well plates at 7 × 10^4^ cells/well and induced overnight with 100 ng/mL doxycycline. Cells were harvested in 2× SDS Laemmli lysis buffer, sonicated, boiled at 100 °C for 10–15 min, and then resolved on a 4–15% Tris-Glycine gel (Bio-Rad). Proteins were transferred to nitrocellulose or PVDF membrane and probed with rat anti-FLAG (clone 9H1, produced in-house; 1:1000) primary antibody and anti-rat HRP (Southern Biotech; 1:5000) secondary.

### TurboID proximity labelling and mass spectrometry analysis

HEL cells (30 × 10^6^ cells) were seeded into T75 flasks and induced overnight with 100 ng/mL doxycycline. The following morning, cells were treated with Biotin (500 µM) for 10 mins and washed 6 times with ice-cold PBS by centrifugation (1500 rpm, 5mins. 4°C) to remove excess biotin. Cells were then lysed in buffer (50 mM Tris-HCl pH 7.4, 1% (v/v) Triton X-100, 1 mM PMSF, 1 mM EDTA, 150 mM NaCl, 2 mM Sodium Vanadate, 10 mM NaF) containing a cOmplete protease inhibitor tablet (Roche) and insoluble debris were removed by centrifugation. Biotinylated proteins were captured from lysates (500 µg per replicate) by incubation with 10 μg high-capacity Streptavidin agarose (Thermo #20359) for 1 hour, rotating at 4°C. The beads were then washed three times with lysis buffer, and three times with PBS + 0.5% SDS before being incubated with 100 µl PBS + 0.5% SDS + 1 mM DTT for 30 minutes at room temperature. The beads were then washed once in 50 mM ammonium bicarbonate + 6 M urea and transferred to Snap Cap Spin Columns (Pierce #69725). Using centrifugation at 1000 X g for 1 min, the beads were washed 5 times 50 mM ammonium bicarbonate + 6 M urea, 5 times with PBS, and 3 times with H_2_O. Proteins were then digested on-bead for 16 hours, at 37°C using 1 µg trypsin (Sigma #EMS0004) in 50 mM ammonium bicarbonate. Peptides were then collected into new tubes by centrifugation. The collected peptides were lyophilized to dryness using a CentriVap (Labconco), before reconstituting in 30 µl 0.1% v/v formic acid/2% v/v acetonitrile ready for mass spectrometry analysis. Peptides were separated by reverse-phase chromatography on a C18 fused silica column (inner diameter 75 µm, OD 360 µm × 15 cm length, 1.6 µm C18 beads) packed into an emitter tip (IonOpticks, Fitzroy, Victoria, Australia), using a nano-flow HPLC (M-class, Waters, Milford, Massachusetts, USA) coupled to a timsTOF Pro (Bruker, Billerica, Massachusetts, USA) equipped with a CaptiveSpray source. Peptides were loaded directly onto the column at a constant flow rate of 400 nl/min with buffer A (99.9% Milli-Q water, 0.1% FA) and eluted with a 30-min linear gradient from 2 to 34% buffer B (99.9% ACN, 0.1% FA). The timsTOF Pro (Bruker) was operated in diaPASEF mode using Compass Hystar 5.1. The settings on the TIMS analyzer were as follows: Lock Duty Cycle to 100% with equal accumulation and ramp times of 100 ms, and 1/K0 Start 0.6 V.·/cm2 End 1.6 V·s/cm2, Capillary Voltage 1400V, Dry Gas 3 l/min, Dry Temp 180°C. The DIA methods were set up using the instrument firmware (timsTOF control 2.0.18.0) for data-independent isolation of multiple precursor windows within a single TIMS scan. The method included two windows in each diaPASEF scan, with window placement overlapping the diagonal scan line for doubly and triply charged peptides in the m/z – ion mobility plane across 16 × 25 m/z precursor isolation windows (resulting in 32 windows) defined from m/z 400 to 1,200, with 1 Da overlap, and CID collision energy ramped stepwise from 20 eV at 0.8 V·s/cm2 to 59eV at 1.3 V·s/cm2.Data files were analysed by DIA-NN v1.8.1 software^48^. Data were searched against the human Uniprot Reference Proteome with isoforms (downloaded May 2021), with recombinant protein sequences added, as a FASTA digest for library-free search with a strict trypsin specificity allowing up to 2 missed cleavages. The peptide length range was set to 7-30 amino acids. Precursor charge range was set between 1-4, and m/z range of 300-1800. Carbamidomethylation of Cys was set as a fixed modification. Precursor FDR was set to 1% and match between runs was enabled. Data processing and analysis of the DIA-NN output were performed using R software (v. 4.2.1). Protein groups were filtered based on a precursor-level q-value of <1% and a protein group-level q-value of <1%, ensuring that only high-confidence identifications were retained. Additionally, only proteins identified with proteotypic peptides were considered for downstream analysis. The PG.MaxLFQ values were used to derive normalised protein group abundances. To ensure data quality, protein groups present in at least 50% of replicates in at least one experimental condition were retained for further analysis. This filtering resulted in a final set of 5,062 proteins. Protein intensity values were subsequently log2-transformed to meet the assumptions of downstream statistical tests. Normalization was carried out using RUVIII-C method (v1.0.19) to reduce unwanted technical variation across samples. Missing values were imputed by drawing random numbers from a normal distribution (with a width parameter of 0.3 and a downshift of 1.8. Principal Component Analysis (PCA) was performed to reduce the dimensionality of the data and identify potential outliers. Differential expression analysis was conducted using the limma R package (v.3.50.0), applying empirical Bayes moderation to improve statistical power. Proteins were deemed significantly differentially expressed if they passed a false discovery rate (FDR) threshold of ≤ 5% after Benjamini– Hochberg (BH) correction. Data visualization was conducted using the ggplot2 R package.

### BiCAP

BiCAP analysis of the PSKH1 dimer was performed as previously described^49, 50^. Briefly, 1×10^6^ HEK293T cells were seeded into 10 cm dishes in 10 mL of growth media 24 h prior to transfection. Samples were prepared in quadruplicate. The two BiCAP expression constructs, pDEST-V1-PSKH1 and pDEST-V2-PSKH2, or a control plasmid expressing full length Venus, were transfected with Jetprime Transfection Agent (Polyplus). Lysates were harvested ∼20 hours after transfection following visual observation of fluorescence. Affinity purification was performed using GFP-Trap® Magnetic Agarose (ChromoTek) beads and captured proteins were prepared for mass spectrometry analysis using the FASP (filter-aided sample preparation) method ^51^, with the following modifications. Proteins were eluted from GFP-Trap beads using 100 μl 0.5% w/v SDS in PBS at 60°C for 3 minutes. Proteins were reduced with 10 mM Tris-(2-carboxyethyl) phosphine (TCEP), alkylated with 50 mM iodoacetamide, then digested with 1 μg sequence-grade modified trypsin gold (Promega) in 50 mM ammonium bicarbonate and incubated overnight at 37°C. Peptides were eluted with 50 mM ammonium bicarbonate in two 40 μl sequential washes and acidified in 1% formic acid (FA, final concentration). The collected peptides were lyophilized to dryness using a CentriVap (Labconco), before reconstituting in 10 µl 0.1% v/v formic acid/2% v/v acetonitrile ready for mass spectrometry analysis. Peptides (3 µl) were separated by reverse-phase chromatography on a C18 fused silica column (inner diameter 75 µm, OD 360 µm × 15 cm length, 1.6 µm C18 beads) packed into an emitter tip (IonOpticks, Austalia) using a custom nano-flow HPLC system (Thermo Ultimate 3000 RSLC Nano-LC, PAL systems CTC autosampler). The HPLC was coupled to a timsTOF Pro (Bruker) equipped with a CaptiveSpray source. Peptides were loaded directly onto the column at a constant flow rate of 400 nL/min in diaPASEF mode as described above. Data files were analysed by DIA-NN v1.8.1 software. Data was searched against the human Uniprot Reference Proteome with isoforms (downloaded August 2022), with recombinant protein sequences added, as a FASTA digest for library-free search with a strict trypsin specificity allowing up to 2 missed cleavages. The peptide length range was set to 7-30 amino acids. Precursor charge range was set between 1-4, and m/z range of 300-1800. Carbamidomethylation of Cys was set as a fixed modification. Precursor FDR was set to 1% and match between runs was on. Data processing and analysis of the DIA-NN output were performed using R software (v. 4.2.1). Protein groups were filtered based on a precursor-level q-value of <1% and a protein group-level q-value of <1%, ensuring that only high-confidence identifications were retained. Additionally, only proteins identified with proteotypic peptides were considered for downstream analysis. The PG.MaxLFQ values were used to derive normalised protein group abundances. To ensure data quality, protein groups present in at least 50% of replicates in at least one experimental condition were retained for further analysis. This filtering resulted in a final set of 4,092 proteins. Protein intensity values were subsequently log_2_-transformed to meet the assumptions of downstream statistical tests. Normalisation was carried out using the cyclic loess method implemented in the limma R package (v. 3.52.2) to reduce unwanted technical variation across samples. Principal Component Analysis (PCA) was performed to reduce the dimensionality of the data and identify potential outliers. Missing values were imputed using the Barycenter (“v2-mnar”) method from the msImpute R package (v. 1.7.0). Differential expression analysis was conducted using the limma R package, applying empirical Bayes moderation to improve statistical power. Proteins were deemed significantly differentially expressed if they passed a false discovery rate (FDR) threshold of ≤ 5% after Benjamini– Hochberg (BH) correction. Data visualisation was conducted using the ggplot2 R package.

### FLAG-IP and mass spectrometry of human PSKH1-3C-FLAG

HEK293T cells were cultured in Dulbecco’s modified Eagle medium (Lonza) supplemented with 10% fetal bovine serum (HyClone), penicillin (50 U/ml), and streptomycin (0.25 µg/ml) (Lonza) and maintained at 37°C in 5% CO2 humidified atmosphere. PSKH1 was cloned into a pcDNA3 backbone vector and expressed in frame with a C-terminal, 3C protease-cleavable FLAG epitope tag. HEK-293T cells were transfected with either PSKH1-3C-FLAG or pCDNA3 EGFP as a control using a 3:1 polyethylenimine (PEI [branched average *M*w ∼25 000 Da; Sigma–Aldrich]) to DNA ratio (30 : 10 µg, for a single 10 cm culture dish). For immunoprecipitation (IP) experiments (all performed in triplicate), proteins were harvested 48 h post transfection in a lysis buffer containing 50 mM Tris–HCl (pH 7.4), 150 mM NaCl, 0.1% (v/v) Triton X-100, 1 mM DTT, 1% (w/v) dodecyl-β-D-Maltoside (DDM), 1 mM ethylenediaminetetraacetic acid (EDTA), 1 mM ethylene glycol-bis(β-aminoethyl ether)-*N*,*N*,*N*′,*N*′-tetraacetic acid (EGTA) and 5% (v/v) glycerol and supplemented with a protease inhibitor cocktail tablet and a phosphatase inhibitor tablet (Roche). Lysates were briefly sonicated on ice, clarified by centrifugation at 20 817×*g* for 20 min at 4°C, and the resulting supernatants were incubated with anti-FLAG G1 Affinity Resin (GenScript) for 3 h with gentle agitation at 4°C. Affinity beads containing bound protein were collected and washed three times in 50 mM Tris–HCl (pH 7.4) and 150 mM NaCl and then equilibrated in storage buffer (50 mM Tris–HCl [pH 7.4], 100 mM NaCl, 1 mM DTT, 1% (w/v) DDM and 5% (v/v) glycerol). The purified proteins were then eluted from the suspended beads over a 3 h period with 3C protease (0.5 µg) at 4°C, with gentle agitation.

IP preparations were diluted to 180 μl in 100 mM ammonium bicarbonate pH 8.0 and reduced and alkylated with dithiothreitol and iodoacetamide, as previously described^52^. Samples were subject to SP3-bead digestion as described^53^ with the adaptions of: using 100 mM ammonium bicarbonate pH 8.0, 0.5 μg of Trypsin gold (Promega) and post-digest bead washing in 1% (w/v) Rapigest (Waters). Once supernatants were combined, samples were acidified by addition of a final concentration of 0.5% TFA and incubated for 30 min each at 37°C with 600 rpm shaking and on ice. Samples were centrifuged at 13,000 *g* for 10 min at 4°C and cleared supernatant collected into fresh tubes prior to vacuum centrifugation. Dried peptides were solubilized in 20 μl of 3% (v/v) acetonitrile and 0.1% (v/v) TFA in water, sonicated for 10 minutes, and centrifuged at 13,000 *g* for 15 min at 4°C prior to reversed-phase HPLC separation using an Ultimate3000 nano system (Dionex) over a 60-minute gradient, as described^52^. All data acquisition was performed using a Thermo QExactive mass spectrometer (Thermo Scientific), with higher-energy C-trap dissociation (HCD) fragmentation set at 30% normalized collision energy for 2+ to 4+ charge states. MS1 spectra were acquired in the Orbitrap (70K resolution at 200 *m/z*) over a *m/z* range of 300 to 2000, AGC target = 1e^6^, maximum injection time = 250 ms, with an intensity threshold for fragmentation of 1e^3^. MS2 spectra were acquired in the Orbitrap (17,500 resolution at 200 *m/z*), maximum injection time = 50 ms, AGC target = 1e^5^ with a 20 s dynamic exclusion window applied with a 10 ppm tolerance. Data was analysed using Proteome Discoverer 2.4 (Thermo Scientific) in conjunction with MASCOT^54^; searching the UniProt Human Reviewed database (updated weekly, accessed September 2024) with constant modifications = carbamidomethyl (C), variable modifications = oxidation (M), instrument type = electrospray ionization–Fourier-transform ion cyclotron resonance (ESI-FTICR), MS1 mass tolerance = 10 ppm, MS2 mass tolerance = 0.01 Da. Analysis was performed as a single study with label free quantification performed using the Minora feature detector node, calculating the area under the curve for *m/z* values, total protein abundance was determined using the HI3 method^55^. Imputation of missing values was performed using low abundance resampling. Data was exported as an Excel file and imported into a custom R script to calculate fold change and t-test statistics for plotting.

### Peptide Array

Positional scanning peptide array experiments and analyses were performed as reported previously^21^. Briefly, recombinant PSKH1 was added to a 384-well plate containing 50 μM peptide substrate library mixtures (Anaspec, AS-62017-1 and AS-62335). The reaction was initiated with the addition of 50 μM ATP (50 μCi ml^−1^ γ-^32^P-ATP, Perkin-Elmer) and incubated for 90 min at 30°C in 50 mM HEPES pH 7.4, 10 mM MgCl_2_, in the presence or absence of 0.2 μM Calmodulin, 400 μM CaCl_2_ (SignalChem catalog # C02-39B-500 calcium/Calmodulin solution); no differences were evident between conditions. Solutions were spotted onto streptavidin-conjugated membranes (Promega, V2861) via the C-terminal biotin on the peptides upon completion of the reaction, before membrane rinsing and imaging using a Typhoon FLA 7000 phosphorimager (GE), and raw data quantification using ImageQuant (GE) to generate densitometry matrices. Densitometry matrices were column-normalized at all positions by the sum of the 17 randomized amino acids (excluding serine, threonine and cysteine), to obtain position-specific scoring matrices (PSSMs).

### In Vitro Kinase Assays

PSKH1 activity was determined by measuring the transfer of radiolabelled phosphate from [γ- ^32^P]-ATP to a synthetic peptide substrate (ADR1; LKKLTRRASFSGQ; synthesized by GenScript, New Jersey, USA). Briefly, purified recombinant GFP-PSKH1 (10 ng) or 10 μL of PSKH1 immobilized on HA-agarose beads (50% v/v) were incubated in assay buffer (50 mM HEPES, pH 7.4, 1 mM DTT) containing 200 μM ADR1 peptide, 200 μM [^32^P]-γ-ATP (Perkin Elmer, MA, USA), 5 mM MgCl_2_ (Sigma) in a 30 μL assay for 10 min at 30°C. Reactions were also supplemented with 100 μM CaCl_2_, 1 μM recombinant CaM (produced in-house) and/ or 1 μM recombinant RCN3 or UNC119B (produced in-house). For the RCN1/RCN3 assays, 1 mM CaCl_2_ and 25 mM MgCl_2_ were used. Reactions were terminated by spotting 15 μL onto phosphocellulose paper (SVI-P; SVI, Melbourne, Australia) and washing extensively in 1% phosphoric acid (Sigma) and dried. Radioactivity was quantified by liquid scintillation counting.

### Chemical crosslinking sample preparation

For SDA crosslinking, recombinant full-length PSKH1 and either recombinant CaM, RCN3 or UNC119B (1:2 molar ratio) were initially incubated on ice in 50 mM HEPES (pH 7.5), 1 mM CaCl_2_ (absent for UNC119B assay) for 30 min. Protein (1 mg/mL) was then mixed with SDA (NHS-Diazirine; 1 mg/mL; Thermo Fisher #26167) and incubated in the dark for 30 min at room temperature to react the NHS-ester group. The diazarine group was then photo-activated using ultraviolet light irradiation (UVP CL-1000L UV cross-linker) at 365 nm. Samples were added to upturned lids excised from 1.5 mL microfuge tubes and placed on ice at a distance of 5 cm from the lamp and irradiated for 1 min. The reaction mixtures from the titration were combined and quenched with 100 mM Tris-HCl (pH 7.5), mixed with reducing sample buffer, heated to 100°C for 5 min, then resolved by reducing SDS-PAGE gel electrophoresis (Bio-Rad). SDS-PAGE gels were stained using SimplyBlue SafeStain (Thermo Fisher) and crosslinked adducts excised for mass spectrometry analysis.

### Chemical crosslinking sample preparation for mass spectrometry-based proteomics

Protein gel bands were excised (based on their migration relative to the MW marker) and destained using 50% acetonitrile for 30 min and 37°C. The gel band was then dehydrated by adding 100% acetonitrile (ACN) and incubating at room temperature for 10 min, before aspirating the ACN and further drying gel piece with the vacuum centrifuge (CentriVap, Labconco). Proteins were reduced by the addition of 1 mM dithiothreitol (DTT) in 50 mM ammonium bicarbonate for 30 min. Excess DTT was aspirated before the addition of 55 mM iodoacetamide to alkylate the sample for 60 min at room temperature. The gel slice was then washed with 50% acetonitrile twice and 100% ACN once before drying to completion in the vacuum centrifuge. Proteins were digested overnight with 500 ng trypsin in 50 mM ammonium bicarbonate at 37°C and extracted the following day using 60% acetonitrile/0.1% formic acid. The collected peptides were lyophilized to dryness using a CentriVap (Labconco), before reconstituting in 10 µL 0.1% formic acid/2% ACN ready for mass spectrometry analysis.

### Chemical crosslinking mass spectrometry analysis

Reconstituted peptides were analyzed on Orbitrap Eclipse Tribrid mass spectrometer interfaced with Neo Vanquish liquid chromatography system. Samples were loaded onto a C18 fused silica column (inner diameter 75 µm, OD 360 µm × 15 cm length, 1.6 µm C18 beads) packed into an emitter tip (IonOpticks) using pressure-controlled loading with a maximum pressure of 1,500 bar using Easy nLC source and electro sprayed directly into the mass spectrometer. We first employed a linear gradient of 3-30% of solvent-B at 400 nL/min flow rate (solvent-B: 99.9% v/v ACN) for 100 min, followed by a gradient of 30-40% solvent-B for 20 min and 35-99% solvent-B for 5 min. The column was then maintained at 99% B for 10 min before being washed with 3% solvent-B for another 10 min comprising a total of 145 min run with a 120 min gradient in a data dependent (DDA) mode. MS1 spectra were acquired in the Orbitrap (R = 120k; normalised AGC target = standard; MaxIT = Auto; RF Lens = 30%; scan range = 380–1400; profile data). Dynamic exclusion was employed for 30s excluding all charge states for a given precursor. Data dependent MS2 spectra were collected in the Orbitrap for precursors with charge states 3-8 (R = 50k; HCD collision energy mode = assisted; normalized HCD collision energies = 25%, 30%; scan range mode = normal; normalised AGC target = 200%; MaxIT = 150 ms). MGF files were searched against a fasta file containing the PSKH1 and interactor sequences using XiSearch software^56^ (version 1.7.6.7) with the following settings: crosslinker = multiple, SDA and noncovalent; fixed modifications = Carbamidomethylation (C); variable modifications = oxidation (M), SDA-loop (KSTY) DELTAMASS:82.04186484, SDA-hydro (KSTY) DELTAMASS:100.052430; MS1 tolerance = 6.0ppm, MS2 tolerance = 20.0ppm; losses = H_2_O, NH_3_, CH_3_SOH, CleavableCrossLinkerPeptide:MASS:82.04186484). FDR was performed with the in-built xiFDR set to 5%. Data were visualized using the XiView software^57^.

### Computational Modelling of PSKH1

AlphaFold model of PSKH1 was obtained from the AlphaFold Protein Structure Database (EMBL-EBI). Richardson (Ribbons) and surface diagrams were drawn using UCSF Chimera.

### Mass spectrometry data deposition

TurboID, BiCAP and FLAG-IP interactome mass spectrometry proteomics data have been deposited to the ProteomeXchange Consortium via the PRIDE^58^ partner repository with the dataset identifiers PXD056543 and PXD055768. SDA crosslinking mass spectrometry data have been deposited to the ProteomeXchange Consortium via jPOST^59^ with accession numbers: JPST003399 and PXD056468 for PSKH1:Calmodulin; JPST003400 and PXD056476 for PSKH1:UNC119B; and JPST003401 and PXD056478 for PSKH1:RCN3.

## Acknowledgments

We thank James Knox and the MiMo crew for facilitating vital discussions; Ben Turk for expert discussions; Professor Naoto Yonezawa for kindly providing the full length RCN1 cDNA; and Professors Alice Ting and Peter Mace for sharing TurboID DNA templates. We thank the National Health and Medical Research Council of Australia for grant (JMM, 1172929; JWS, 2001817) and infrastructure (9000719) support; the Australian Research Council for grant support (JWS, DP210102840); the Victorian State Government Operational Infrastructure Support Scheme; and UKRI BBSRC grant BB/S018514/1 for funding (PAE, CEE, DPB, LAD).

## Author Contributions

CRH, JWS and JMM conceptualized the study and wrote the manuscript; CRH, TAD, SNY, LJM, JLJ, LAD, DPB, ALC, BHN, DHM, LS, LMMc, JWS designed and performed experiments, and analysed data; TMY-B, EMH, JY, GM, ARM analysed data; CRH, LFD, LCC, MCT, DRC, CEE, PAE, JWS, JMM supervised the study and provided resources; all authors edited the manuscript.

---

**Figure S1.**
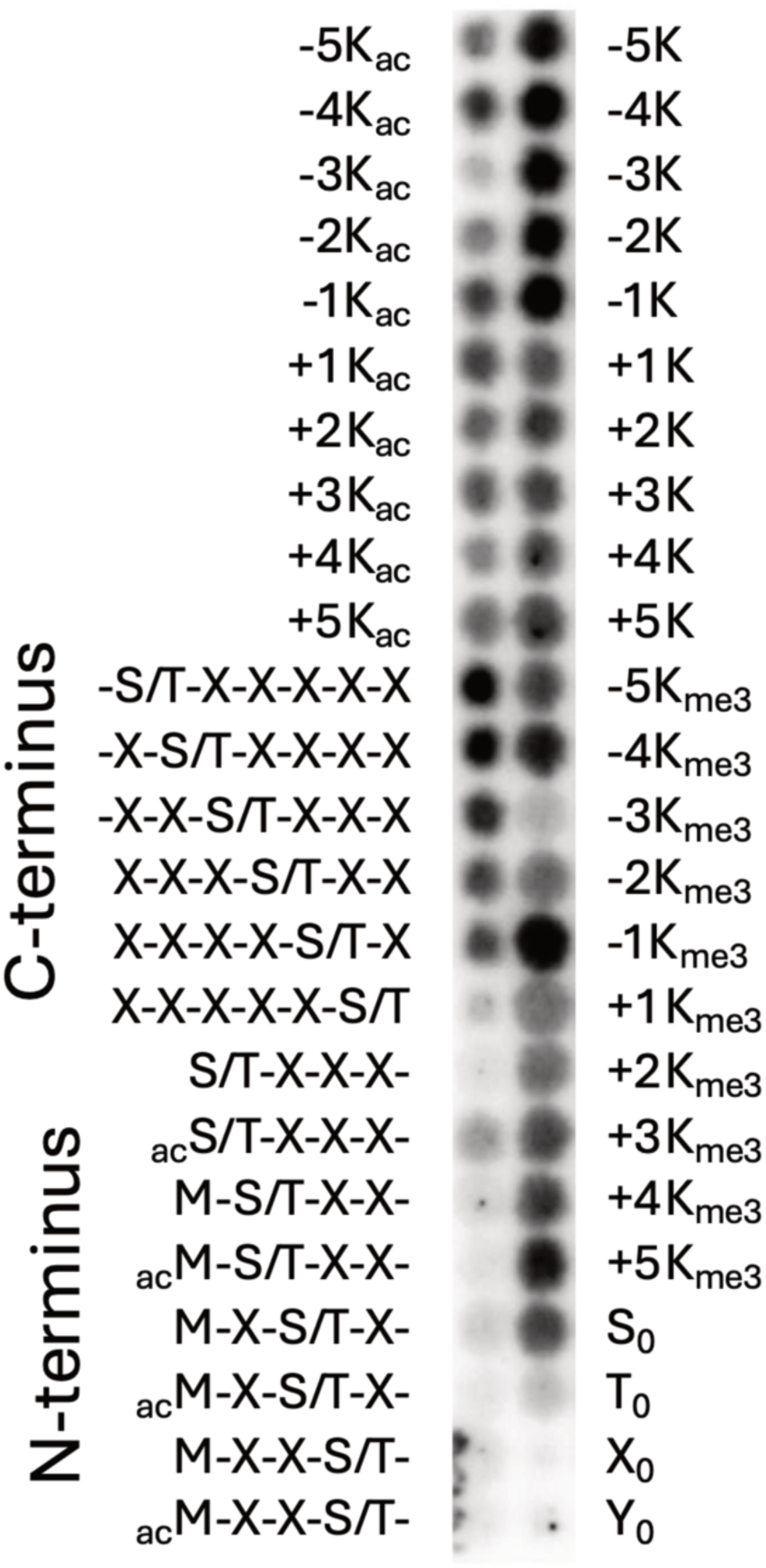
PSKH1 prefers basic residues at the −3 position and favors C-terminal phosphoacceptors. Lysine (K) residues are well tolerated at all positions, although loss of charge by acetylation (Ac) or trimethylation (Me3) at the −3 position compromises recognition by PSKH1. S_0_, G-A-X-X-X-X-X-S-X-X-X-X-A-G-K-K(LC-biotin); T_0_, G-A-X-X-X-X-X-T-X-X-X-X-A-G-K-K(LC-biotin); X_0_, G-A-X-X-X-X-X-X-X-X-X-X-A-G-K-K(LC-biotin); Y_0_, G-A-X-X-X-X-X-Y-X-X-X-X-A-G-K-K(LC-biotin); where X = degenerate mixture of the 16 natural amino acids excluding cysteine, tyrosine, serine, and threonine.

## References

1. Berginski ME, Moret N, Liu C, Goldfarb D, Sorger PK, Gomez SM. The Dark Kinase Knowledgebase: an online compendium of knowledge and experimental results of understudied kinases. Nucleic Acids Res 49, D529–D535 (2021).

2. Whitworth H, et al. Identification of kinases regulating prostate cancer cell growth using an RNAi phenotypic screen. PLoS One 7, e38950 (2012).

3. Maddirevula S, et al. Large Scale Genomic Investigation of Pediatric Cholestasis Reveals a Novel Hepatorenal Ciliopathy Caused by PSKH1 Mutations. Genet Med, 101231 (2024).

4. Brede G, Solheim J, Stang E, Prydz H. Mutants of the protein serine kinase PSKH1 disassemble the Golgi apparatus. Exp Cell Res 291, 299–312 (2003).

5. Horne CR, et al. Unconventional binding of Calmodulin to CHK2 kinase inhibits catalytic activity. bioRxiv, (2024).

6. Brede G, Solheim J, Troen G, Prydz H. Characterization of PSKH1, a novel human protein serine kinase with centrosomal, golgi, and nuclear localization. Genomics 70, 82–92 (2000).

7. Chin D, Means AR. Calmodulin: a prototypical calcium sensor. Trends Cell Biol 10, 322–328 (2000).

8. Kahl CR, Means AR. Regulation of cell cycle progression by calcium/calmodulin-dependent pathways. Endocr Rev 24, 719–736 (2003).

9. Babu YS, Sack JS, Greenhough TJ, Bugg CE, Means AR, Cook WJ. Three-dimensional structure of calmodulin. Nature 315, 37–40 (1985).

10. Zhang M, Tanaka T, Ikura M. Calcium-induced conformational transition revealed by the solution structure of apo calmodulin. Nat Struct Biol 2, 758–767 (1995).

11. Honore B. The rapidly expanding CREC protein family: members, localization, function, and role in disease. Bioessays 31, 262–277 (2009).

12. Peersen OB, Madsen TS, Falke JJ. Intermolecular tuning of calmodulin by target peptides and proteins: differential effects on Ca2+ binding and implications for kinase activation. Protein Sci 6, 794–807 (1997).

13. Suzuki N, et al. Calcium-dependent structural changes in human reticulocalbin-1. J Biochem 155, 281–293 (2014).

14. Mazzorana M, Hussain R, Sorensen T. Ca-Dependent Folding of Human Calumenin. PLoS One 11, e0151547 (2016).

15. Wright KJ, et al. An ARL3-UNC119-RP2 GTPase cycle targets myristoylated NPHP3 to the primary cilium. Genes Dev 25, 2347–2360 (2011).

16. Cardenas-Rodriguez M, et al. Genetic compensation for cilia defects in cep290 mutants by upregulation of cilia-associated small GTPases. J Cell Sci 134, (2021).

17. Jaiswal M, Fansa EK, Kosling SK, Mejuch T, Waldmann H, Wittinghofer A. Novel Biochemical and Structural Insights into the Interaction of Myristoylated Cargo with Unc119 Protein and Their Release by Arl2/3. J Biol Chem 291, 20766–20778 (2016).

18. Cen O, Gorska MM, Stafford SJ, Sur S, Alam R. Identification of UNC119 as a novel activator of SRC-type tyrosine kinases. J Biol Chem 278, 8837–8845 (2003).

19. Hutti JE, et al. A rapid method for determining protein kinase phosphorylation specificity. Nat Methods 1, 27–29 (2004).

20. Songyang Z, Blechner S, Hoagland N, Hoekstra MF, Piwnica-Worms H, Cantley LC. Use of an oriented peptide library to determine the optimal substrates of protein kinases. Curr Biol 4, 973–982 (1994).

21. Johnson JL, et al. An atlas of substrate specificities for the human serine/threonine kinome. Nature 613, 759–766 (2023).

22. Yaron-Barir TM, et al. The intrinsic substrate specificity of the human tyrosine kinome. Nature 629, 1174–1181 (2024).

23. Chen C, et al. Identification of a major determinant for serine-threonine kinase phosphoacceptor specificity. Mol Cell 53, 140–147 (2014).

24. Branon TC, et al. Efficient proximity labeling in living cells and organisms with TurboID. Nat Biotechnol 36, 880–887 (2018).

25. Uhlen M, et al. Proteomics. Tissue-based map of the human proteome. Science 347, 1260419 (2015).

26. Byrne DP, et al. Evolutionary and cellular analysis of the ‘dark’ pseudokinase PSKH2. Biochem J 480, 141–160 (2023).

27. Taipale M, et al. Quantitative analysis of HSP90-client interactions reveals principles of substrate recognition. Cell 150, 987–1001 (2012).

28. Tachikui H, Navet AF, Ozawa M. Identification of the Ca(2+)-binding domains in reticulocalbin, an endoplasmic reticulum resident Ca(2+)-binding protein with multiple EF-hand motifs. J Biochem 121, 145–149 (1997).

29. de Diego I, Kuper J, Bakalova N, Kursula P, Wilmanns M. Molecular basis of the death-associated protein kinase-calcium/calmodulin regulator complex. Sci Signal 3, ra6 (2010).

30. Komolov KE, et al. Structure of a GRK5-Calmodulin Complex Reveals Molecular Mechanism of GRK Activation and Substrate Targeting. Mol Cell 81, 323–339 e311 (2021).

31. Simon B, et al. Death-Associated Protein Kinase Activity Is Regulated by Coupled Calcium/Calmodulin Binding to Two Distinct Sites. Structure 24, 851–861 (2016).

32. Constantine R, Zhang H, Gerstner CD, Frederick JM, Baehr W. Uncoordinated (UNC)119: coordinating the trafficking of myristoylated proteins. Vision Res 75, 26–32 (2012).

33. Mejuch T, van Hattum H, Triola G, Jaiswal M, Waldmann H. Specificity of Lipoprotein Chaperones for the Characteristic Lipidated Structural Motifs of their Cognate Lipoproteins. Chembiochem 16, 2460–2465 (2015).

34. Yelland T, Garcia E, Samarakoon Y, Ismail S. The Structural and Biochemical Characterization of UNC119B Cargo Binding and Release Mechanisms. Biochemistry 60, 1952–1963 (2021).

35. Shrestha S, Byrne DP, Harris JA, Kannan N, Eyers PA. Cataloguing the dead: breathing new life into pseudokinase research. FEBS J 287, 4150–4169 (2020).

36. Brede G, Solheim J, Prydz H. PSKH1, a novel splice factor compartment-associated serine kinase. Nucleic Acids Res 30, 5301–5309 (2002).

37. Li Y, et al. Global genetic analysis in mice unveils central role for cilia in congenital heart disease. Nature 521, 520–524 (2015).

38. Honore B, Vorum H. The CREC family, a novel family of multiple EF-hand, low-affinity Ca(2+)-binding proteins localised to the secretory pathway of mammalian cells. FEBS Lett 466, 11–18 (2000).

39. Vorum H, Liu X, Madsen P, Rasmussen HH, Honore B. Molecular cloning of a cDNA encoding human calumenin, expression in Escherichia coli and analysis of its Ca2+-binding activity. Biochim Biophys Acta 1386, 121–131 (1998).

40. Ismail SA, Chen YX, Miertzschke M, Vetter IR, Koerner C, Wittinghofer A. Structural basis for Arl3-specific release of myristoylated ciliary cargo from UNC119. EMBO J 31, 4085–4094 (2012).

41. Stephen LA, Ismail S. Shuttling and sorting lipid-modified cargo into the cilia. Biochem Soc Trans 44, 1273–1280 (2016).

42. Huttlin EL, et al. The BioPlex Network: A Systematic Exploration of the Human Interactome. Cell 162, 425–440 (2015).

43. Huttlin EL, et al. Architecture of the human interactome defines protein communities and disease networks. Nature 545, 505–509 (2017).

44. Huttlin EL, et al. Dual proteome-scale networks reveal cell-specific remodeling of the human interactome. Cell 184, 3022–3040 e3028 (2021).

45. Fitzgibbon C, Meng Y, Murphy JM. Co-expression of recombinant RIPK3:MLKL complexes using the baculovirus-insect cell system. Methods Enzymol 667, 183–227 (2022).

46. Meng Y, et al. Human RIPK3 C-lobe phosphorylation is essential for necroptotic signaling. Cell Death Dis 13, 565 (2022).

47. Murphy JM, et al. The pseudokinase MLKL mediates necroptosis via a molecular switch mechanism. Immunity 39, 443–453 (2013).

48. Demichev V, Messner CB, Vernardis SI, Lilley KS, Ralser M. DIA-NN: neural networks and interference correction enable deep proteome coverage in high throughput. Nat Methods 17, 41–44 (2020).

49. Croucher DR, et al. Bimolecular complementation affinity purification (BiCAP) reveals dimer-specific protein interactions for ERBB2 dimers. Sci Signal 9, ra69 (2016).

50. Hastings JF, et al. Dissecting Multi-protein Signaling Complexes by Bimolecular Complementation Affinity Purification (BiCAP). J Vis Exp, (2018).

51. Wisniewski JR, Zougman A, Nagaraj N, Mann M. Universal sample preparation method for proteome analysis. Nat Methods 6, 359–362 (2009).

52. Ferries S, et al. Evaluation of Parameters for Confident Phosphorylation Site Localization Using an Orbitrap Fusion Tribrid Mass Spectrometer. Journal of Proteome Research 16, 3448–3459 (2017).

53. Daly LA, et al. Custom Workflow for the Confident Identification of Sulfotyrosine-Containing Peptides and Their Discrimination from Phosphopeptides. Journal of Proteome Research 22, 3754–3772 (2023).

54. Perkins DN, Pappin DJ, Creasy DM, Cottrell JS. Probability-based protein identification by searching sequence databases using mass spectrometry data. Electrophoresis 20, 3551–3567 (1999).

55. Silva JC, Gorenstein MV, Li G-Z, Vissers JPC, Geromanos SJ. Absolute Quantification of Proteins by LCMSE: A Virtue of Parallel ms Acquisition *S. Molecular & Cellular Proteomics 5, 144–156 (2006).

56. Mendes ML, et al. An integrated workflow for crosslinking mass spectrometry. Molecular Systems Biology 15, e8994 (2019).

57. Graham M, Combe C, Kolbowski L, Rappsilber J. xiView: A common platform for the downstream analysis of Crosslinking Mass Spectrometry data.). bioRxiv (2019).

58. Perez-Riverol Y, et al. The PRIDE database resources in 2022: a hub for mass spectrometry-based proteomics evidences. Nucleic Acids Res 50, D543–D552 (2022).

59. Okuda S, et al. jPOSTrepo: an international standard data repository for proteomes. Nucleic Acids Res 45, D1107–D1111 (2017).

